# THLANet: A Deep Learning Framework for Predicting TCR-pHLA Binding in Immunotherapy Applications

**DOI:** 10.1101/2025.04.14.648678

**Authors:** Xu Long, Qiang Yang, Weihe Dong, Xiaokun Li, Kuanquan Wang, Suyu Dong, Gongning Luo, Xianyu Zhang, Tiansong Yang, Xin Gao, Guohua Wang

**Affiliations:** School of Computer Science and Technology, Harbin Institute of Technology, Harbin, 150001, China; School of Computer Science and Technology, Heilongjiang University, Xuefu Road, Harbin, 150080, China; College of Computer and Control Engineering, Northeast Forestry University, Harbin, 150004, China; Department of Breast Surgery, Harbin Medical University Cancer Hospital, Harbin 150081, China; Computer Science Program, Computer, Electrical and Mathematical Sciences and Engineering Division, King Abdullah University of Science and Technology (KAUST), Thuwal 23955-6900, Kingdom of Saudi Arabia; Center of Excellence for Smart Health (KCSH), King Abdullah University of Science and Technology (KAUST), Thuwal 23955-6900, Kingdom of Saudi Arabia; Department of Rehabilitation, The First Affiliated Hospital of Heilongjiang University of Traditional Chinese Medicine, Xuefu Road, 150040, Harbin, China; Postdoctoral Program of Heilongjiang Hengxun Technology Co., Ltd., Xuefu Road, 150090, Harbin, China; Shandong Hengxun Technology Co., Ltd., Miaoling Road, 266100, Qingdao, China

## Abstract

Adaptive immunity is a targeted immune response that enables the body to identify and eliminate foreign pathogens, playing a critical role in the anti-tumor immune response. Tumor cell expression of antigens forms the foundation for inducing this adaptive response. However, the human leukocyte antigens (HLA)-restricted recognition of antigens by T-cell receptors (TCR) limits their ability to detect all neoantigens, with only a small subset capable of activating T-cells. Accurately predicting neoantigen binding to TCR is, therefore, crucial for assessing their immunogenic potential in clinical settings. We present THLANet, a deep learning model designed to predict the binding specificity of TCR to neoantigens presented by class I HLAs. THLANet employs evolutionary scale modeling-2 (ESM-2), replacing the traditional embedding methods to enhance sequence feature representation. Using scTCR-seq data, we obtained the TCR immune repertoire and constructed a TCR-pHLA binding database to validate THLANet’s clinical potential. The model’s performance was further evaluated using clinical cancer data across various cancer types. Additionally, by analyzing divided complementarity-determining region (CDR3) sequences and simulating alanine scanning of antigen sequences, we unveiled the 3D binding conformations of TCRs and antigens. Predicting TCR-neoantigen pairing remains a significant challenge in immunology, THLANet provides accurate predictions using only the TCR sequence (CDR3*β*), antigen sequence, and class I HLA, offering novel insights into TCR-antigen interactions.

**Author summary:** T-cell receptor (TCR) recognition of peptide-human leukocyte antigen (pHLA) complexes is fundamental to immune responses. However, predicting their binding poses a significant challenge due to the intricate dynamics of their interactions. We developed THLANet, a novel deep learning model, to address this challenge by integrating the ESM-2 and Transformer-Encoder modules. This approach enhances sequence feature encoding, improving the model’s generalization capability and enabling accurate predictions of TCR-pHLA binding in clinical datasets. Using data processed from open-source databases, THLANet outperformed existing methods, such as PanPep and pMTnet, in precision-recall metrics across multiple epitopes. Additionally, THLANet demonstrates superior capability in identifying critical binding sites within 3D structures, providing structural insights into TCR-pHLA interactions. THLANet offers a robust framework for advancing immunotherapy research, with potential applications in the development of personalized medicine.

## Introduction

Cancer arises from genetic aberrations, including small variants like single-nucleotide substitutions, insertions, and deletions [1–3]. These alterations are unique to cancer cells and absent in normal tissues [4, 5]. As a result, protein products derived from these mutations hold the potential to deliver clinical benefits without inducing toxicity in healthy tissues [6–8]. The emergence advent of strategies that activate harness the host immune system to combat malignant tumors cells represents marks a significant paradigm pivotal shift in the field of cancer immunotherapy [9–12]. Recently, immunotherapies such as adoptive cell therapy (ACT) [13, 14], Immune checkpoint blockade, and tumor antigen vaccines have achieved significant clinical success [13, 20, 21]. The therapeutic efficacy largely stems from the anti-tumor activity of CD8+ T-cells, which target malignant cells by recognizing tumor-associated neoantigens and tumor-specific antigens on their surface [15–17]. Consequently, accurately evaluating a T-cell repertoire’s capacity to interact with tumor antigens is essential for identifying and targeting cancer cells and remains central to optimizing cancer immunotherapy [18, 19, 22].

Human leukocyte antigen (HLA) molecules present neoantigenic peptides that are recognized by T-cell receptors (TCRs), triggering the transformation of naive T-cells into CD8+ cytotoxic T-cells. This activation stimulates the immune system, enabling the targeted destruction of malignant cells (Fig. 1a) [23]. Each T-cell clone possesses a unique TCR, which can act as an antigenic peptide to safeguard the immune system against malignancies [24]. The complementarity-determining region 3 (CDR3) of the TCR *β*-chain is particularly significant due to its highly diverse antigen specificity, making it the focal point of our study on TCR-neoantigen interactions. Given that TCRs must account for HLA constraints when recognizing neoantigens—acknowledging both antigenic peptides and polymorphisms—understanding the binding mechanism of the TCR ternary complex is essential. The ternary complex, comprising the TCR, antigenic peptide, and HLA (TCR-pHLA), is crucial for autoimmune antigen discovery and cancer vaccine development. Advances in artificial intelligence computational models have enabled precise prediction and identification of TCR-pHLA pairings, paving the way for groundbreaking research in modern immunology [25].

**Fig 1.**
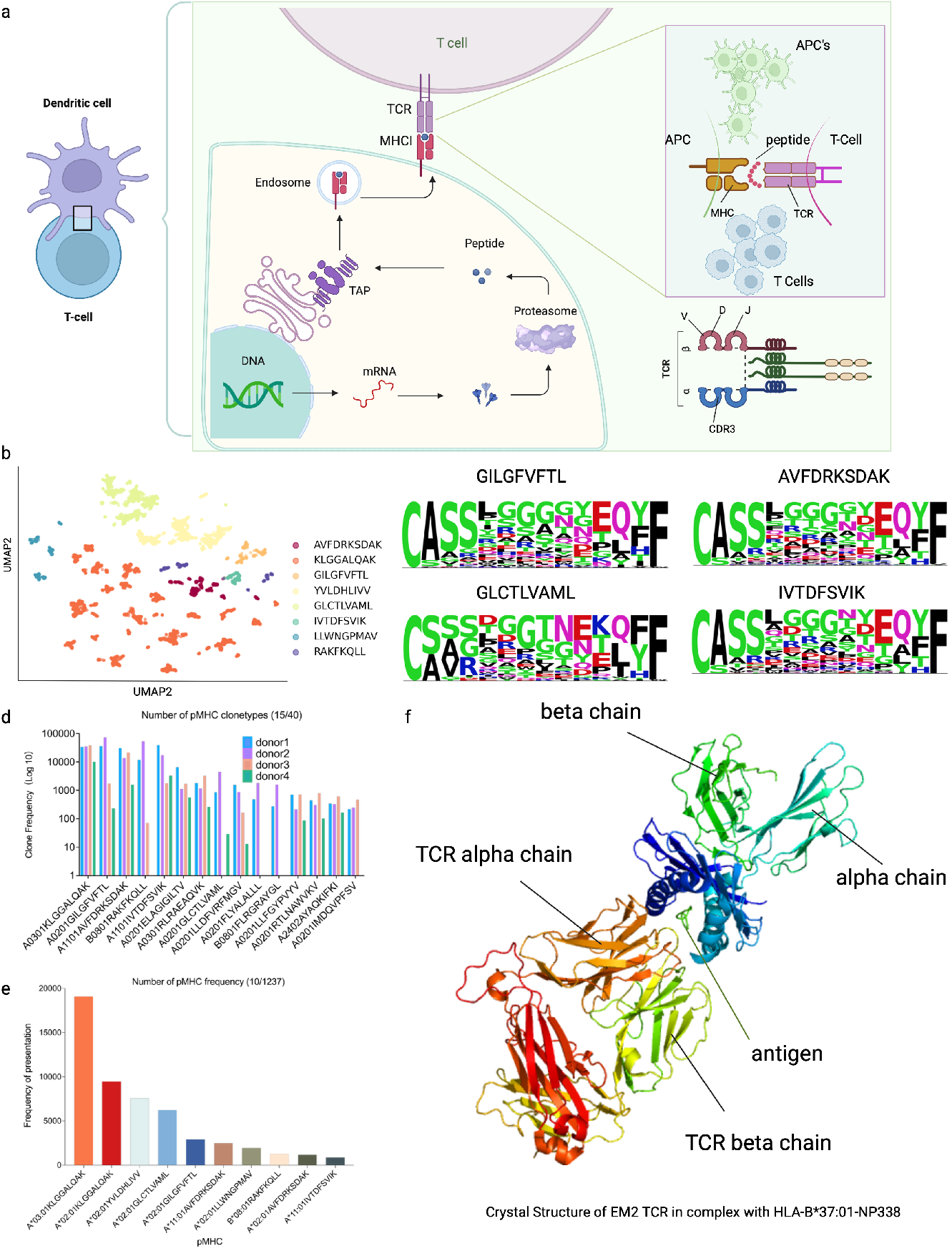
Statistical information on immunogenicity data. (**a**) Schematic diagram of TCR–peptide-major histocompatibility complex (pMHC) recognition by CD8+ T-cells. (**b**) Uniform manifold approximation and projection (UMAP) projections of the predicted epitope-specific TCR clusters. (**c**) Sequence motifs of CDR3*β* representing the epitope-specific TCRs for the four protein epitopes. (**d**) Distribution of pMHC clonality in data from four donors using 10X Genomics. (**e**) Distribution of pMHC data from the VDJ database (VDJdb) and immune epitope database (IEDB). (**f** ) 3D binding schematic of TCR-pHLA (from PDB ID: 6MTM).

**Fig 2.**
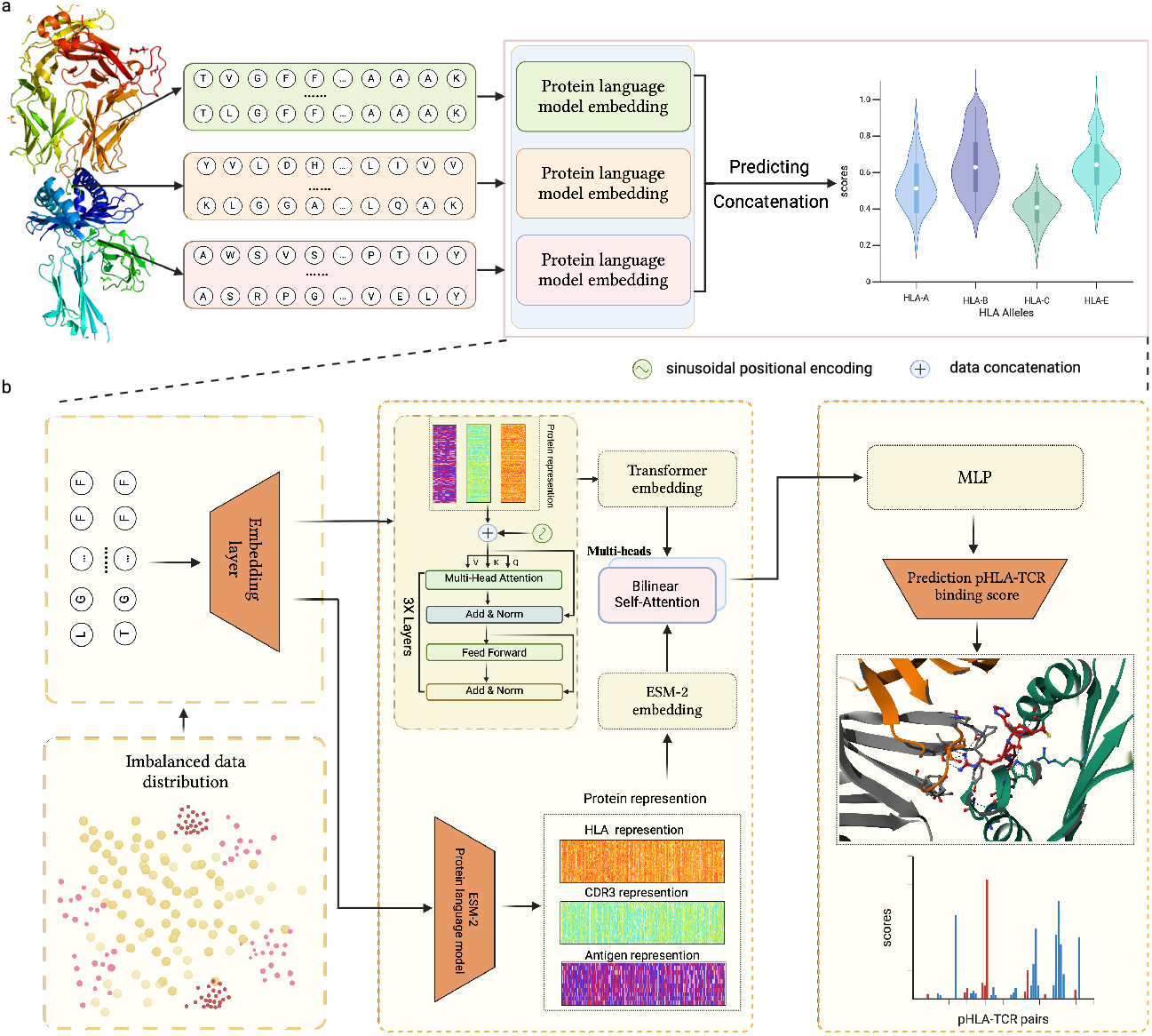
Workflow of THLANet for predicting T-cell receptor (TCR)-peptide-human leukocyte antigen (pHLA) interactions. (**a**) The pipeline of THLANet for predicting the TCR-pHLA triad group interaction process. (**b**) Detailed architecture of THLANet: The training data exhibit a long-tail distribution. Protein sequences are processed through the ESM2 module and a convolutional neural network (CNN) to capture long-distance dependencies and features. Concurrently, the sequences are initially encoded using the BLOSUM62 matrix and embedded through a transformer encoder. The two feature matrices are fused via a bilinear attention network. In the prediction module, a multilayer perceptron (MLP)-based model predicts the interaction scores between TCR and pHLA.

Several computational tools are available to analyze TCR models and predict antigenic peptide-TCR binding specificity. Previous studies have classified these tools into three categories: (1) tools for defining TCR clusters and deciphering antigen-specific binding models, such as TCRdist [26], DeepTCR [27], GIANA [28], and GLIPH2 [29]; (2) tools for predicting antigen-peptide specific TCR binding models, including T-cell receptor gaussian process (TCRGP) [30], TCRex [31], and NetTCR-2.0 [32]; and (3) tools for developing pan-peptide TCR binding prediction models trained on known bound TCRs, such as pMTnet [33], PanPep [34], TARB-BERT [35], DLpTCR [36], and TITAN [37]. However, the first and second categories lack the robustness needed to address the broad-spectrum TCR-pHLA recognition problem. Furthermore, models in the third category have not sufficiently explored the TCR-pHLA binding mechanism.

We propose THLANet, a deep learning framework leveraging evolutionary scale modeling (ESM-2) embeddings to predict the binding specificity between class I pHLA and TCR [38]. To validate the model, we incorporated multiple independent datasets, including THLANet’s assessment of the TCR-pHLA 3D binding conformation, providing insights into the 3D binding mechanism [39–42]. Additionally, we compiled cancer-related TCR-pHLA binding data from various peer-reviewed publications and applied THLANet for predictions, demonstrating its potential for clinical applications in cancer [43, 44].

## Materials and methods

### Data curation

In this study, we curated multiple datasets to support our analysis, including a foundational dataset, a 3D crystal structure dataset of TCR-pHLA complexes, and a dataset of tumor-mutant immunogenic antigens.

We began by analyzing data generated using the 10x Genomics Chromium single-cell immunoassay platform, which leverages feature barcoding technology to create single-cell 5’ libraries and V(D)J-enriched TCR sequence libraries. This platform also employs highly multiplexed pHLA multimers to identify binding specificity. Specifically, we examined four single-cell datasets obtained from the peripheral blood of four healthy donors with no known viral infections (As shown in Section E in S1 Text). These datasets included 44 CD8+ T-cell profiles that demonstrated specific binding to pHLA complexes. We investigated the clonal expansions of T cells to identify TCRs capable of interacting with pMHC complexes, quantifying them using original unique molecular identifiers (UMIs). When the UMI count reached or exceeded 10, the pMHC-TCR pair was deemed capable of interacting with T cells and was classified as a positive sample. From these single-cell immunoassay datasets, we compiled TCR-pHLA datasets containing 19,657 entries.

To ensure data quality, we processed the data using the open-source databases VDJdb (N = 36,168) and the IEDB (N = 31,836), resulting in high-confidence datasets [39, 40]. Given the critical role of the CDR3 region of the TCR beta chain in epitope recognition, we focused exclusively on recording the CDR3 beta chain. The integrated dataset (N = 87,661) was further refined by removing duplicates and dichotomous samples, yielding a final dataset of 66,314 unique entries. We retained laboratory-validated negative samples from the IEDB database and selected samples with a confidence score of 0 from the VDJdb database, collectively constructing the negative dataset for our study.

As shown in Supplementary Fig. 4 in S1 Text and Fig. 1d , we analyzed cell type distributions across the four single-cell sequencing datasets and examined the clonotype distribution of pHLA. A total of 40 pHLA sequence datasets demonstrated significant clonal expansions (*>*10 clones) in donor T-cells. We visualized the clonal distribution of the top 15 pHLAs in CD8+ T-cells (Fig. 1d). Additionally, we cataloged 1,237 distinct pHLA types across all datasets, evaluated their overall frequencies, and highlighted the top 10 pHLA types by occurrence (Fig. 1e). The pronounced coupling of certain pHLA types underscores their frequent occurrence in clinical settings, validating the clinical relevance of our findings.

For the TCR-pHLA crystal structure data, we curated 3D structural information from screenings in the IEDB and retrieved 198 crystal structure datasets from the PDB database using their ID information [41, 42]. After excluding low-quality entries, we finalized a dataset of 112 high-quality TCR-pHLA crystal structures. The spatial configuration of the pHLA–TCR triad is illustrated in Figure 1f (derived from PDB ID: 6MTM).

We also explored the physiological significance of TCR-pHLA interactions characterized by THLANet, focusing on tumor neoantigens arising from somatic mutations. To this end, we compiled TCR-pHLA pairs from peer-reviewed studies involving tumor-infiltrating lymphocytes derived from melanoma and gastrointestinal cancers [43, 44]. These pairs were identified through high-throughput immunology screenings, enabling the characterization of TCR-pHLA interactions in these two highly immunogenic tumor types. This analysis offers clinically relevant insights into the mechanisms of tumor antigen presentation within the tumor microenvironment.

### Data Embedding

We employed evolutionary scale modeling 2 (ESM-2) for protein sequences to encode amino acid sequences [38]. ESM-2 is an unsupervised learning language model based on the transformer architecture, trained with up to 15 billion parameters. Utilizing an attention mechanism, it identifies interaction patterns between amino acid pairs in the input sequences. ESM-2 plays a pivotal role in ESMFold’s success in predicting protein structures. The deep features embedded by ESM-2 capture critical information about the three-dimensional conformation of protein structures, offering theoretical insights into the binding mechanisms of TCR-pHLA interactions in three-dimensional space. As a pretrained model, ESM-2 retains a vast repository of prior protein knowledge, which increases the risk of overfitting. To address this and enrich encoded features, we first processed protein sequences using the Blocks Substitution Matrix 62 (Blosum62) and applied sinusoidal encoding to enhance spatial sequence representation [45, 46]. For unified encoding with ESM-2, we utilized the transformer encoder architecture to effectively embed the data. To seamlessly integrate the datasets generated by ESM-2 and the transformer encoder while minimizing information loss or conflicts, we implemented a bilinear attention network. This approach efficiently fuses the two embeddings, preserving the rich joint features and enabling a comprehensive and accurate representation.

As shown in Figure 1b, we performed a two-dimensional uniform manifold approximation and projection (UMAP) analysis of antigen-associated samples processed by the bilinear attention network. The samples, representing the top eight antigens by frequency in the training dataset, were visualized to assess the encoded feature distribution. The visualization clearly demonstrates that the encoded data exhibits distinct distribution patterns, with samples grouped by antigen type displaying well-defined regional clustering within the two-dimensional space.

### Bilinear attention network

The bilinear attention network is an attention-based model designed primarily for multimodal learning tasks, such as visual question answering (VQA) [47]. In VQA, given an image and a natural language question, the system generates an answer by matching text and image data. The bilinear attention network excels in preserving rich joint information while maintaining computational efficiency, offering a promising approach for enabling interaction between ESM-2 and transformer encoder data in our study [48].

### Pseudosequence Construction and Data Preparation

The pseudosequence of the HLA sequence is constructed by selecting 34 amino acid residues to characterize the polymorphism of HLA-A, -B, and -C alleles, with these residues identified as the positions within the binding groove most likely to directly interact with the antigenic peptide, thereby effectively capturing the interaction characteristics between the antigenic peptide and the HLA molecule. The selection of these 34 amino acids is based on their critical role in various binding conformations, ensuring a robust representation of HLA-peptide interactions [49]. To prepare the data for modeling, the input sequences are aligned by standardizing the lengths of antigenic residues and CDR3*β* to 15 and 25, respectively, with shorter sequences padded using the placeholder *<*pad*>* to ensure uniformity. Additionally, the original dataset is partitioned into training, validation, and test sets in an 8:1:1 ratio, respectively, to facilitate subsequent model training and evaluation.

### THLANet architecture

#### ESM-2 architecture

ESM-2, the most advanced protein language model to date, incorporates significant advancements in architecture, training parameters, computational resources, and data scale. Trained on protein sequences from the UniRef database, ESM-2 employs a masked prediction approach where 15% of amino acids are randomly masked, and the model is tasked with predicting these positions. This training objective emphasizes amino acid prediction while requiring the model to capture complex internal representations of input sequences to achieve high accuracy. ESM-2 encodes each protein sequence into a 1280-dimensional feature vector, enabling detailed exploration of protein sequence characteristics and serving as a robust tool for protein research. For input processing, ESM-2 prepends a *<*cls*>* token at the beginning of each sequence to mark its start and appends an *<*eos*>* token at the end to denote its conclusion. When applied to encode antigenic peptides, HLA sequences, and CDR3*β* regions, the resulting feature vectors are concatenated to form a unified matrix with the dimensions [n, 79, 1280], where n represents the number of samples, 79 corresponds to the total length of the concatenated sequence (including the standardized lengths and special tokens), and 1280 is the dimensionality of the feature vector for each position.

#### Transformer-Encoder architecture

To enhance the feature richness of the prediction samples, we employed the Transformer-Encoder module to generate a novel encoding matrix. Each pHLA-TCR pair was encoded into a 79×21 matrix, referred to as the TCR-pHLA matrix, using the BLOSUM62 matrix. To incorporate positional information, sinusoidal positional encoding was applied to each amino acid within the matrix. This encoding method enables the model to capture the relative distances and sequential relationships between amino acid positions within the protein, thereby guiding the model’s attention mechanism more effectively based on positional context. The resulting TCR-pHLA matrix, now augmented with positional information, was utilized as the input to the Transformer encoder. After processing through three layers of the Transformer encoder, the final encoded matrix was obtained, in which each amino acid is enriched with global contextual information derived from the entire sequence.

### THLANet prediction model

The THLANet prediction model integrates of CNN layers with MLP layers. The CNN (TextCNN) employs three 1D convolutional layers with kernel sizes of [1, 2, 1] and 1000 filters to extract features from text sequences. ReLU activation and max pooling operations follow the convolutional layers, and the pooled outputs are concatenated into a feature vector. This vector passes through three fully connected layers with 1048 and 512 units, each activated by ReLU, followed by batch normalization layers to stabilize training. A dropout layer (rate = 0.4) mitigates overfitting. Finally, a softmax layer outputs a binary classification probability distribution. Model optimization utilized the Adam optimizer with a learning rate of 5 * 10^*−*4^ and a batch size of 128. Training ran for up to 100 epochs, with early stopping triggered if validation loss failed to decrease for five consecutive epochs.

### CDR*β* has binding preference

We analyzed the amino acid preferences of CDR3*β* sequences associated with responses to eight distinct protein epitopes: AVFDRKSDAK, KLGGALQAK, GILGFVFTL, YVLDHLIVV, GLCTLVAML, IVTDFSVIK, LLWNGPMAV, and RAKFKQLL (Fig. 1c and Supplementary Fig. 2 in S1 Text). While the N- and C-terminal regions of the epitope-specific CDR3*αβ* loops exhibited relative conservation, the regions directly interacting with the epitopes showed substantial diversity. Interestingly, specific amino acids such as glycine (G), leucine (L), proline (P), and polar residues like asparagine (N), serine (S), threonine (T), and tyrosine (Y) were more frequently identified in the core positions of CDR3*β* sequences.

### Experimental setting

#### THLANet training

THLANet was implemented using Python 3.10, PyTorch 2.5.0, and Compute Unified Device Architecture (CUDA) 12.4. The computational environment included an Intel(R) Xeon(R) Gold 6248R CPU @ 3.00GHz with 256 GB of RAM and two NVIDIA A100-PCIE-40GB GPUs, each offering 40 GB of memory. The Adam optimizer minimized the binary cross-entropy loss. The model achieved complete convergence after 60 epochs.

#### Performance evaluation metrics

The performance was assessed using AUROC, AUPR. In AUROC (TPR versus FPR for a series of threshold values) 1, the true-positive rate (TPR) and false-positive rate (FPR) are computed as:

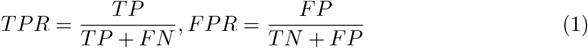

Here TP denotes true positive; FN, false negative; TN, true negative; and FP, false positive. In AUPR 2, the precision and recall are computed by:

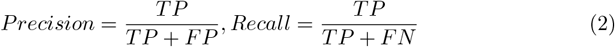

### Statistical analyses

Performance benchmarking metrics, including the area under the curve (ROC-AUC) and area under the precision-recall curve (PR-AUC), were computed using the Python package scikit-learn v.1.3.0. UMAP was conducted with the Python package umap-learn v.0.5.5. Sequence motifs were visualized using the Python package clogmaker v.0.8 with the “weblogo_protein” color scheme [50]. PyMOL v.2.5.8 was employed to visualize the 3D structure of TCR–pMHC complexes.

## Result

### THLANet Outstanding Performance

In this study, we evaluated THLANet, pMTNet, PanPep, and TARB-BERT using a test set from THLANet, comprising samples from commonly used open-source databases and clinically relevant TCR-pHLA samples derived through rigorous data cleaning of the 10X Genomics Datasets, these models are described in Section F in S1 Text. This approach enabled each model to predict scenarios closely resembling clinical conditions. Benchmarking against existing tools, as illustrated in Fig. 3a, revealed that THLANet outperformed the other three baseline models. THLANet achieved superior performance with median receiver operating characteristic (ROC)-AUC and precision-recall (PR)-AUC scores of 0.8803 and 0.8552, respectively. In comparison, pMTNet scored 0.8664 (ROC-AUC) and 0.7883 (PR-AUC), while PanPep performed worse with scores of 0.8384 and 0.7231. TARB-BERT displayed intermediate results, achieving median ROC-AUC and PR-AUC scores of 0.8686 and 0.8405. Notably, PanPep and pMTNet exhibited poor performance in ROC-AUC and PR-AUC metrics. TARB-BERT, however, delivered better results, likely due to incorporating the BERT protein pretraining model. This allowed for high-dimensional spatial encoding of short peptide proteins, enabling the model’s ability to learn features effectively. These findings underscore the potential of protein pretraining models to leverage data characteristics, thereby advancing the application of antigen recognition in clinical settings.

**Fig 3.**
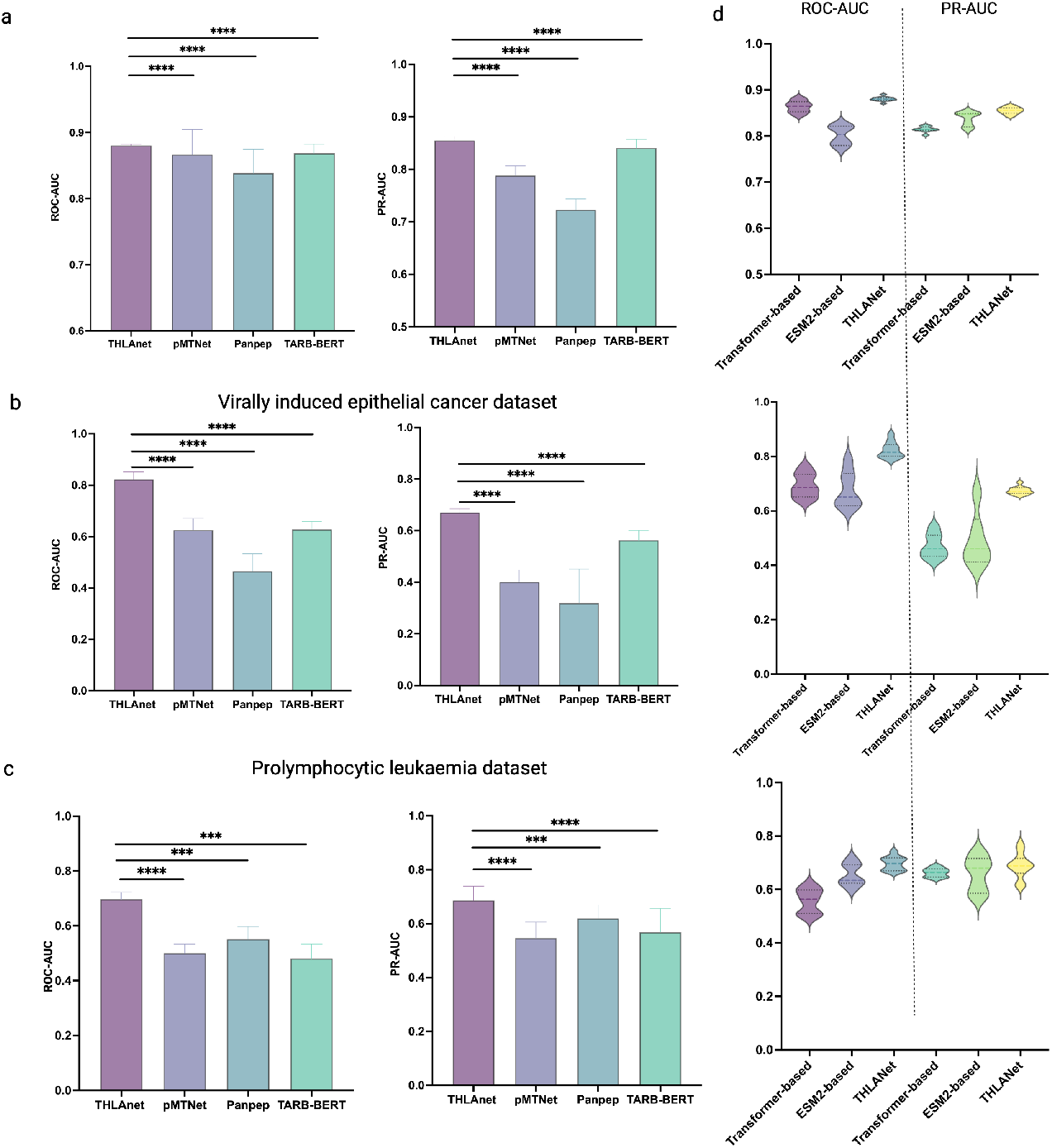
THLANet prediction results. (**a**) Receiver operating characteristic–area under the curve (ROC-AUC) and precision-recall–area under the curve (PR-AUC) in the testing dataset. (**b**) ROC-AUC and PR-AUC in elanoma and gastrointestinal cancer datasets. (**c**) ROC-AUC and PR-AUC in prolymphocytic leukemia dataset. (**d**) Ablation experiment , ROC-AUC and PR-AUC with two-tailed Wilcoxon signed-rank test adjusted *p*-values across methods.

We evaluated the AUPR of epitopes for representative TCRs in the test dataset. THLANet emerged as the best predictor for 12 of the 19 epitope targets (Fig. 4), including the EBV-BRLF1 epitope YVLDHLIVV (AUPR: 0.9883), the yellow fever virus peptide LLWNGPMAV (AUPR: 0.9045), and the hepatitis C virus peptide ATDALMTGY (AUPR: 0.9611).

**Fig 4.**
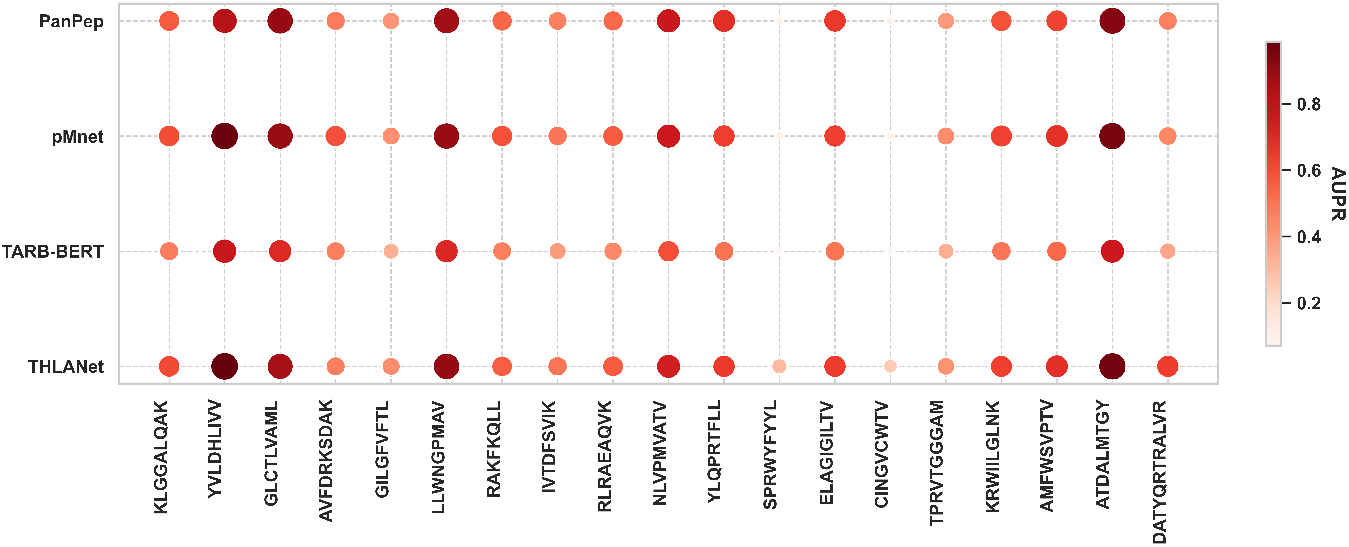
Comparison of area under the precision-recall curve (AUPR) values derived from PanPep, pMNet, TARB-BERT, and THLANet for 19 epitopes with more than ten binding T-cell receptors (TCRs) in the test dataset. A darker color and larger point size indicate a higher AUPR.

### THLANet exhibits potential in multi-cancer clinical applications

The interaction between TCRs and neoantigens is crucial in driving the progress of tumor immunotherapy. To assess whether THLANet can efficiently identify and predict neoantigens across various cancer types in clinical settings and validate its potential as a knowledge discovery tool, we characterized TCR-pHLA (T-cell receptor–human leukocyte antigen) interactions in several highly immunogenic tumor types.

We analyzed data from a patient with T-cell prolymphocytic leukemia (T-PLL) provided by 10X Genomics. This dataset utilizes feature barcode technology to generate single-cell 5 libraries and V(D)J-enriched libraries for TCR sequencing, along with highly multiplexed pHLA multimer reagents to determine binding specificity. The dataset includes 6,365 TCR clonotypes responsive to T-PLL, with samples having unique molecular identifiers (UMIs) ≥10 classified as positive and those with UMIs *<* 10 classified as negative. Furthermore, we incorporated peer-reviewed data from melanoma and gastrointestinal cancers, yielding a total of 6,365 TCR-pHLA pairs associated with T-PLL and 54 pairs linked to melanoma and gastrointestinal cancers [43, 44] (Supplementary Table S5 and S6 in S1 Text).

Using THLANet, we predicted TCR-pHLA interactions in clinical cancer samples and compared its performance against three baseline models. In predictions for melanoma and gastrointestinal cancer samples, as expected, THLANet outperformed pMTNet, PanPep, and TARB-BERT, achieving median ROC-AUC and PR-AUC scores of 0.8152 and 0.6845, respectively (compared to pMTNet’s median ROC-AUC and PR-AUC scores of 0.625 and 0.4088, respectively; PanPep’s 0.5200 and 0.3416; and TARB-BERT’s 0.6391 and 0.5744) (Fig. 3b).

Similarly, in predictions for T-PLL samples, THLANet achieved superior performance, with a median ROC-AUC of 0.6976 and a median PR-AUC of 0.7170 (compared to pMTNet’s median ROC-AUC of 0.5055 and PR-AUC of 0.5713; PanPep’s median ROC-AUC of 0.5445 and PR-AUC of 0.5949; and TARB-BERT’s median ROC-AUC of 0.4800 and PR-AUC of 0.5635) (Fig. 3c).

These findings underscore THLANet’s ability to predict peptide-specific TCR binding interactions across diverse cancer types effectively, highlighting its significant potential for advancing exogenous TCR binding studies.

In conclusion, our study reaffirmed the practicality of THLANet in identifying neoantigens that elicit immune responses in CD8+ T-cells, highlighting its significant potential in advancing immune therapies such as ACT, TCR-T, and CAR-T strategies.

### Ablation experiment

We investigated the impact of the two primary modules in the network architecture on the performance of THLANet, focusing on the transformer encoder and ESM-2 architectures. To conduct this analysis, ablation studies were performed on each architecture individually, followed by a comparative evaluation using the validation dataset and two peer-reviewed cancer-related TCR-pHLA datasets (Fig. 3d). The study revealed that both the transformer encoder and ESM-2 architectures moderately improved. While the ESM-2 architecture demonstrated exceptional ability in identifying positive samples, it showed less stability in performance compared to the transformer encoder. By integrating both architectures, THLANet was able to leverage their respective strengths, achieving significant advancements in both predictive performance and stability.

We retrained and evaluated the model under the condition that only the Transformer encoder was retained as the encoding module. We observed the following results on the test dataset: AUROC increased by 0.00173, and AUPRC increased by 0.0373. While the improvement in AUROC was not statistically significant at the 0.05 significance level (two-tailed Wilcoxon signed-rank test), the increase in AUPRC was significant (*p <* 0.05). On two independent cancer-related datasets, the transformer encoder achieved improvements of AUROC = 0.1302 (*p* = 0.0078) and AUPRC = 0.1444 (*p <* 0.05), as well as AUROC = 0.1394 (*p <* 0.05) and AUPRC = 0.0309 (*p <* 0.05).

Under the ablation of the Transformer encoder module, we retrained and evaluated the model. Leveraging the pretrained protein parameters of the ESM-2 model, we embedded protein sequences to capture richer encoding features. Following ablation of the ESM-2 architecture, we retrained and reevaluated the model. On the test dataset, we observed AUROC increased by 0.00804, and AUROC increase of 0.0139 (*p <* 0.05). On two independent cancer-related datasets, the ESM-2 architecture achieved improvements of AUROC = 0.2038, AUPRC = 0.1830, and AUROC = 0.0258 (*p <* 0.005), AUPRC = 0.0309 (*p <* 0.005).

According to the findings of this experiment, the integration of the ESM-2 module and the Transformer-Encoder module provides richer sequence encoding features for THLANet, while also enhancing the overall generalization capability of the model, enabling more accurate predictions of clinical sample data.

### THLANet provides explanation for the spatial structure of the TCR-pHLA interaction

To investigate the binding mechanisms of TCR-pHLA interactions in the microenvironment of CD8+ T-cells, we performed computational simulations of mutation analyses to identify significant changes in TCR and pHLA binding caused by mutations in CDR3 residues. To support our experiments and validate our findings in 3D space, we compiled 112 samples from the IEDB containing 3D structures of TCR-pHLA complexes, the data filtering criteria are provided in Section D of S1 Text. The alanine scanning technique used in biophysical studies provided the conceptual framework for this analysis. Using computational simulations of alanine scanning, we performed base mutations on all TCRs in the 112 test groups, recording the predicted differences between the wild-type TCRs and their mutated counterparts.We divided each TCR CDR3 region into five equal-length segments (Fig. 5a). As expected, residues in the central segment of CDR3 induced significantly greater variations in predicted binding affinity compared to those in the external segments (first, second, and fifth segments) (t-test, *p <* 0.0001 when comparing the third or fourth segment with any of the remaining segments). In Fig 5a–c, we present the TCR-pHLA structure generated by Daichao Wu using X-ray diffraction (PDB ID: 8GON, Resolution: 2.60 Å). Following computational simulations of alanine scanning, residue-by-residue predictions were conducted (Fig. 5b). The residues SER101 and ARG99 exhibited the most significant predictive differences, attributed to their central locations in CDR3, where they made the most contact with pHLA (Fig 5c). Notably, the SER101 residue simultaneously contacts two antigen residues, resulting in even greater predictive changes. In summary, our study demonstrates that, although THLANet was developed at the one-dimensional sequence level, it exhibits the capacity to elucidate the intrinsic properties of peptide-TCR interactions and to identify critical sites within three-dimensional structures.

**Fig 5.**
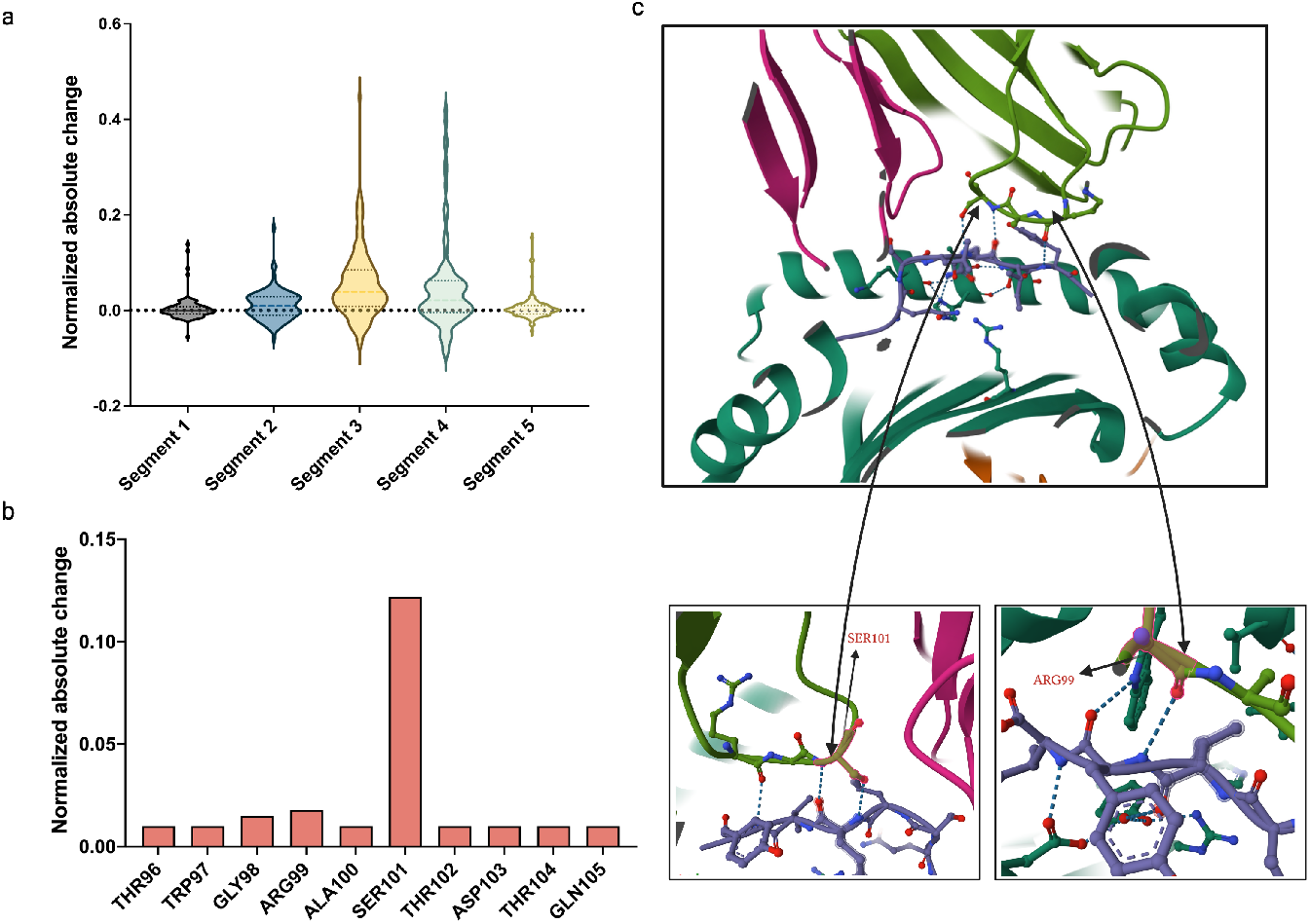
Validation of the T-cell receptor–human leukocyte antigen (TCR-pHLA) network (THLANet) in identifying critical sites within the three-dimensional (3D) crystal structure. (**a**) The complementarity-determining region 3 (CDR3) residues in the middle of the TCR sequence exhibited significant changes in scores predicted by THLANet, as validated through alanine scanning mutagenesis. The CDR3 sequence was divided into five equal-length segments for the alanine scanning analysis. (**b-c**) Predicted score changes for amino acid residues in the CDR3 of an example TCR–peptide-major histocompatibility complex TCR-pMHC structure (PDB ID: 8GON). The 3D structure of 8GON is illustrated: green, CDR3 of the TCR*β* chain; magenta, TCR*α* chain; tints, other regions of the TCR*β* chain; violet, antigen.

## Discussion

In this study, we introduce THLANet, a novel deep learning framework designed to efficiently predict TCR-neoantigen binding specificity. Since not all neoantigens elicit a T-cell response, the ability to predict which TCRs recognize specific neoantigens is essential for tailoring personalized therapies. Our findings emphasize the importance of accurately identifying TCR-pHLA interactions to optimize cancer immunotherapy. THLANet outperformed existing models across evaluation metrics, demonstrating greater potential for clinical applications. By leveraging the ESM-2 approach, THLANet enhances prediction accuracy by exploiting rich sequence features, providing deeper insights into the binding mechanisms of TCRs to antigens at the three-dimensional structural level. Computational simulations explored and revealed the binding conformation of TCRs and pHLAs, elucidating how specific residues influence binding affinity. This structural understanding is invaluable for guiding future therapeutic strategies and designing more effective cancer vaccines. THLANet shows significant promise, and future work will involve further validation across diverse cancer patient populations to assess its robustness in various cancer types. Additionally, integrating other molecular data may further enhance its predictive capabilities.

In conclusion, THLANet represents a significant advancement in optimizing T-cell-mediated immunotherapy, paving the way for more effective cancer treatments. Several directions for future research include: (1) enhancing THLANet to predict the binding of antigens presented by HLA class II molecules to TCRs and (2) leveraging the synergistic integration of THLANet with genomic technologies to validate predicted neoantigens in immunotherapy across a broader range of cancer types and diverse patient cohorts.

## Supporting information

**S1 Text. Supplementary materials**.

## Author contributions

**Conceptualization**: Xu Long, Xiaokun Li.

**Data curation**: Xu Long, Qiang Yang.

**Formal analysis**: Weihe Dong, Suyu Kong.

**Funding acquisition**: Xiaokun Li, Kuanquan Wang, Gongning Luo.

**Investigation**: Xu Long, Qiang Yang.

**Methodology**: Xu Long.

**Project administration**: Xu Long, Xin Gao, Guohua Wang.

**Supervision**: Xiaokun Li, Xin Gao, Guohua Wang.

**Validation**: Xu Long.

**Writing – original draft**: Xu Long.

**Writing – review & editing**: Xianyu Zhang, Tiansong Yang.

## Acknowledgments

The authors thank the reviewers for their valuable comments and suggestions.

